# Density dependent enhancement effect of *Wolbachia* and the host RNAi response to a densovirus in *Aedes* cells

**DOI:** 10.1101/441584

**Authors:** Rhys Parry, Cameron Bishop, Lachlan de Hayr, Sassan Asgari

## Abstract

The endosymbiotic bacterium *Wolbachia pipientis* has been shown to restrict a range of RNA viruses in *Drosophila melanogaster* and transinfected dengue mosquito, *Aedes aegypti*. Here, we show that *Wolbachia* infection enhances replication of Aedes albopictus densovirus (AalDNV-1), a single stranded DNA virus, in *Aedes* cell lines in a density-dependent manner. Analysis of previously produced small RNAs of Aag2 cells showed that *Wolbachia-*infected cells produced greater proportions of viral derived short interfering RNAs as compared to uninfected cells. Additionally, we found production of viral derived PIWI-like RNAs (vpiRNA) produced in response to AalDNV-1 infection. Nuclear fractions of Aag2 cells produced a primary vpiRNA signature U_1_ bias whereas the typical “ping-pong” signature (U_1_ - A_10_) was evident in the cytoplasmic fraction. This is the first report of the density-dependent enhancement of DNA viruses by *Wolbachia*. Further, we report the generation of vpiRNAs in a DNA virus-host interaction for the first time.

## Introduction

Aedes aegypti densovirus (AeDNV) and Aedes albopictus densovirus (AalDNV) are small non-enveloped, single-stranded DNA viruses belonging to the monosense *Brevidensovirus* genus of the *Parvoviridae* family (Bergoin and Tijssen, 2010). Members of brevidensoviruses have a 4-kb genome with three open reading frames on the same strand encoding for two non-structural proteins (NS1, NS2) and a capsid protein (VP) (Bergoin and Tijssen, 2010), as well as a small ORF with unknown function on the complementary strand (Afanasiev et al., 1991). Unique palindromic sequences of approximately ~140-160 nt at the 3′ and 5′ end of the genome fold to form a T-shaped structure (Afanasiev et al., 1991). Densoviruses replicate and form paracrystalline arrays in the nuclei of mosquito cells and cause characteristic nuclear hypertrophy (densonucleosis) (Carlson et al., 2006).

AeDNV, AalDNV and also Anopheles gambiae densovirus (AgDNV) have been intensively studied as transducing vectors and also as biocontrol agents (Reviewed by (Carlson et al., 2006; Johnson and Rasgon, 2018). Both *Ae. aegypti* and *Ae. albopictus* are vectors of a diverse range of medically relevant viruses from the *Togaviridae* and *Flaviviridae* families such as chikungunya virus (CHIKV), dengue virus (DENV) and Zika virus (ZIKV) (Bhatt et al., 2013; Juliano and Lounibos, 2005; Parry and Asgari, 2018). Aedes densoviruses are attractive biocontrol agents as in-laboratory studies of *Ae. aegypti* and *Ae. albopictus* mosquitoes infected with AeDNV and AalDNV reveal a spectrum of DNV pathological effects (Carlson et al., 2006). In addition, AeDNV and AalDNV have been demonstrated to have limited host range, infectious to all life stages and unable to infect other insect and mammalian hosts (Jousset et al., 1993). AalDNV has been demonstrated to cause up to 97.6% mortality in the first instar *Ae. aegypti* larvae (Barreau et al., 1996). Whereas AeDNV has been shown to cause 66% and 64% larval and pupal mortality in *Ae. aegypti* and *Ae. albopictus*, respectively (Buchatsky et al., 1997). Aedes Thailand densonucleosis virus (AThDNV) was demonstrated to produce a larval mortality rate of 51% and 82% in *Ae. aegypti* and *Ae. albopictus*, respectively (Kittayapong et al., 1999). Further examination of AeDNV-induced mortality in *Ae. aegypti* mosquitoes by Ledermann *et al.* suggested that mortality is a dose-dependent effect with peak mortality of up to 75% (Ledermann et al., 2004).

*Wolbachia pipientis* is a gram-negative endosymbiotic bacterium present in approximately 40–65 % of insect species, diverse arthropods and nematodes (Hilgenboecker et al., 2008; Jeyaprakash and Hoy, 2000). Endosymbiotic *Wolbachia* infections of *Drosophila melanogaster* have been demonstrated to reduce viral loads when challenged with dsRNA and positive sense RNA viruses (Hedges et al., 2008; Teixeira et al., 2008). Despite the ubiquitous presence of *Wolbachia* within insect hosts, a worldwide survey of *Ae. aegypti* from 27 countries and six continents showed no presence of *Wolbachia* (Gloria-Soria et al., 2018). Only a single report of a naturally occurring infection of *Wolbachia* in *Ae. aegypti* mosquitoes has been published which suggests that a natural *Wolbachia* infection in *Ae. aegypti* mosquitoes is absent or exceedingly rare (Coon et al., 2016). In contrast, *Ae. albopictus* mosquitoes are naturally infected with two strains of *Wolbachia* (*w*AlbA and *w*AlbB), which can be single infections or superinfections of both strains (Sinkins et al., 1995; Zhou et al., 1998).

Stable introduction of *Wolbachia w*Mel and *w*MelPop from *D. melanogaster* into *Ae. aegypti* mosquitoes and cell lines has been shown to restrict replication and dissemination of medically important single stranded RNA viruses such as DENV, ZIKV, Yellow fever virus, and West Nile virus from the family *Flaviviridae* (Aliota et al., 2016a; Dutra et al., 2016; Hussain et al., 2013a; Moreira et al., 2009; van den Hurk et al., 2012; Walker et al., 2011; Ye et al., 2015), as well as members of *Togaviridae* CHIKV and Mayaro virus (Aliota et al., 2016b; Moreira et al., 2009; van den Hurk et al., 2012). Stable transinfection of *w*Mel in *Ae. albopictus* has also been demonstrated to restrict DENV and CHIKV in the mosquito host (Blagrove et al., 2012; Blagrove et al., 2013). In addition to stable *w*Mel/*w*MelPop transinfections, it has also been demonstrated that stable infection of *Wolbachia* strain *w*AlbB from *Ae. albopictus* into *Ae. aegypti* mosquitoes increases *Ae. aegypti* DENV refractoriness (Ant et al., 2018; Bian et al., 2010; Joubert and O’Neill, 2017; Joubert et al., 2016).

Efforts to elucidate interactions between *Wolbachia* and the insect host have almost primarily focused on positive sense single stranded RNA viruses. It has been reported that stable transinfection of *w*MelPop-CLA in *Ae. aegypti* cells Aag2 restricts the single stranded positive sense RNA virus Cell Fusing Agent virus (CFAV, *Flaviviridae*) (Schnettler et al., 2016; Zhang et al., 2016) but does not restrict the single stranded negative sense RNA virus Phasi Charoen like virus (PCLV, *Bunyaviridae*) (Schnettler et al., 2016) nor the single stranded negative sense RNA virus Aedes anphevirus (family not assigned) (Parry and Asgari, 2018) within these cells.

Currently, only two reports have characterised the interaction of *Wolbachia*-infected insects with DNA viruses. *D. melanogaster* infected with *w*Mel showed an increased mortality when challenged with Invertebrate iridescent virus 6 (IIV-6), (Family, *Iridoviridae*), compared to the tetracycline cleared fly line (Teixeira et al., 2008). Titration of IIV-6 accumulation in challenged flies showed a non-significant 1.8-fold increase in the presence of *Wolbachia* as compared to uninfected flies (Teixeira et al., 2008). In the second report, the presence of three different *Wolbachia* strains *w*Exe1, *w*Exe2, and *w*Exe3 in the African armyworm moth *Spodoptera exempta* was positively associated with prevalence of Spodoptera exempta nucleopolyhedrovirus (SpexNPV) related deaths in the moth’s larvae (Graham et al., 2012). Laboratory bioassays also demonstrated that the presence of *Wolbachia* increased susceptibility and decreased survival of *S. exempta* challenged with SpexNPV (Graham et al., 2012)

A number of Aedes densoviruses have been isolated from persistently infected mosquito cell lines that show no observable cytopathic effects (Boublik et al., 1994; Chen et al., 2004; O’Neill et al., 1995). To examine the possibility of persistent densovirus infection in our Aag2 cell line and also the possible interactions between *Wolbachia* infection and densovirus, we conducted RNA sequencing and also re-analyzed small RNA profiles of viral RNAs (vRNAs) in Aag2 and Aag2.*w*MelPop-CLA cells produced previously from the cytoplasmic and nuclear fractions (Mayoral et al., 2014a; Mayoral et al., 2014b). In addition, we also explored the effect of a supergroup B *Wolbachia* strain (wAlbB) on Aedes densovirus by generating a stably transinfected Aag2 cell line with *w*AlbB (denoted Aag2-*w*AlbB).

## Methods and Materials

### Mosquito cells

*Ae. aegypti* Aag2 cell line (Peleg, 1968) and Aag2 cells infected with *w*MelPop-CLA, previously described by (Frentiu et al., 2010), *Ae. albopictus* Aa23 cells previously described by (O’Neill et al., 1997) and *Ae. albopictus* C6/36 cells (Singh, 1967) were maintained in 1:1 Mitsuhashi-Maramorosch and Schneider’s insect medium (Invitrogen) supplemented with 5–10% fetal bovine serum (FBS, Bovogen Biologicals, French origin), while Aa20 cells (Pudney et al., 1979) were maintained in L15 medium (Invitrogen) supplemented with 10% tryptose phosphate broth (TPB) and 5% FBS. All mosquito cell lines were kept at 28 °C and passaged every 3–4 days. Tetracycline treatment of cell lines has been described in (Asad et al., 2018).

### Production of stably infected Aag2-*w*AlbB cell line

*Wolbachia w*AlbB strain from *Ae. albopictus* Aa23 cells were transinfected into the Aag2 cell line using a method adapted from (Iturbe-Ormaetxe et al., 2011). Briefly, Aa23 cells were maintained in a 1:1 mixture of Mitsuhashi–Maramorosch and Schneider’s insect media (Invitrogen), supplemented with 10% FBS, in a 175cm2 culture flask until confluent. Cells were then lysed via sonication, and the lysate filtered through a 1.2 μm filter. The filtrate was centrifuged and resuspended in 2 ml of 1:1 Mitsuhashi–Maramorosch and Schneider’s insect media, supplemented with 10% FBS. The suspension was added to 1.5 x 106 Aag2 cells adhered to 1 well of a 6-well plate, and allowed to incubate for 24 h at 27°C. Thereafter, cells were resuspended and maintained in a 1:1 mixture of Mitsuhashi–Maramorosch and Schneider’s insect media, supplemented with 10% FBS, and passaged regularly at high density to increase *Wolbachia* density. Samples were taken at each alternate passage and tested for *Wolbachia* density via qPCR targeting the *Wolbachia* Surface Protein gene (WSP) (forward: 5′-ATCTTTTATAGCTGGTGGTGGT-3′; reverse:5′ GGAGTGATAGGCATATCTTCAAT-3′) and *Aa. aegypti* Ribosomal Protein Subunit 17 gene (RPS17) (forward: 5-CACTCCCAGGTCCGTGGTAT-3′; reverse 5′- GGACACTTCCGGCACGTAGT-3′). To generate a *Wolbachia* free Aag2-*w*AlbB line (Aag2-*w*AlbBT) we tetracycline cured using the methods outlined in (Asad et al., 2018).

### Experimental inoculation of Aa20 cells with AalDNV-1

Aa20 cells were experimentally infected with AalDNV-1 from persistently infected Aag2 and C6/36 cell lines as follows. 6×10^6^ Aa20 cells were seeded into a 12 well plate and allowed to adhere for 30 min. To prepare AalDNV-1 inoculum from cell lines, C6/36 and Aag2 cells were spun at 16000 xg for 3 min. The supernatant was then filtered through a 0.22 μm filter. Aa20 medium was removed and replaced with 1mL of inoculation medium. Cells were rocked for 1 h, the first time point was designated as the first time point day 0. Samples were then taken at 1, 3 and 5 dpi.

### RNA extraction and sequencing

Total RNA of Aag2 cells was extracted using QIAzol Reagent (Qiagen). Total RNA was quantified and qualified by Agilent 2100 bioanalyzer (Agilent Technologies, Palo Alto, CA, USA), NanoDrop (Thermo Fisher Scientific Inc.) and 1% agarose gel. 1 μg total RNA with RIN value above 7 was used for the following library preparation. Next generation sequencing library preparations were constructed according to the manufacturer’s protocol using NEBNext First Strand Synthesis Reaction Buffer and NEBNext Random Primers. First strand cDNA was synthesized using ProtoScript II Reverse Transcriptase and the second-strand cDNA was synthesized using Second Strand Synthesis Enzyme Mix. The purified double-stranded cDNA (by AxyPrep Mag PCR Clean-up, Axygen) was then treated with End Prep Enzyme Mix to repair both ends and add a dA-tailing in one reaction, followed by a TA ligation to add adaptors to both ends. Size selection of adaptor-ligated DNA was then performed using AxyPrep Mag PCR Clean-up, and fragments of ~360 bp (with the approximate insert size of 300 bp) were recovered. Each sample was then amplified by PCR for 11 cycles using P5 and P7 primers, with both primers carrying sequences which can anneal with flow cell to perform bridge PCR and P7 primer carrying a six-base index allowing for multiplexing.

### DNA Extraction and PCR Amplification

DNA was extracted using a DIY spin column protocol described in (Ridley et al., 2016). A primer was designed to amplify a 298 bp region in the NS1 protein for both AalDNV-1 (forward: 5’-AGTGAACATTCGCCGTGTGA-3’; reverse 5’-CTCTGGAGCCGCTGTGTAAT-3’). Three step with melt qPCR analysis was undertaken using QuantiFast SYBR Green PCR Kit (QIAGEN, Germany) and using a Rotor-Gene Q machine (QIAGEN, Germany) with 200ng of input DNA (5μL SYBER Green, qF/qR primer 0.25μL, H_2_O 0.5μL). Amplification was performed at 95 °C for 1 min, followed by 35 cycles of 95 °C for 30 s, 56 °C for 30 s, 68 °C for 1 min and a final extension at 68 °C for 5 min. Control primers for host were the *Ae. aegypti RPS17* gene and also to the *wsp* gene for *Wolbachia.* All PCR products were individually inspected using melt curve analysis, ran and eluted from a 1% agarose gel and Sanger sequenced (AGRF, Brisbane).

### Total and small RNA sequencing analysis

For processing of total RNA-Seq data we used CLC Genomics Workbench (version 10.1.1) to remove adapter sequences and reads with low quality scores from datasets. We applied the quality score of 0.05 as cut off for trimming. As described in CLC Genomic Workbench manual, the program uses the modified-Mott trimming algorithm for this purpose. For assembly of the CDS regions of densovirus from Aag2 cells we used the CLC *de novo* assembler with default word length and bubble sizes. Assembled contigs were then queried using BLASTn against the AalDNV-1 strain (Genbank ID: X74945).

For processing the Aag2 and the Aag2.*w*MelPop-CLA small RNA (sRNA) libraries, we used the approach previously described (Mayoral et al., 2014a). For normalization and comparison between sRNA libraries we used reads per million. Unmapped reads less than 16 nt and greater than 32 nt were trimmed from all libraries and sRNA was then mapped to our assembled contig using the strict mapping criteria (mismatch, insertion and deletion costs: 2 : 3 : 3, respectively). For analysis of nucleotide frequency and conservation of AalDNV-1 piRNAs, the RNA-Seq tool on CLC Genomics Workbench was used with default mapping parameters. Mapped reads were then extracted, trimmed to individual nucleotide lengths and visualized using WebLogo (Crooks et al., 2004).

### Phylogenetic analysis

To determine relatedness between the assembled CDS region of Aag2 densovirus and the previously reported densovirus species of Culicidae mosquitoes, we aligned the CDS regions using the ClustalW algorithm on CLC Genomics Workbench. A Maximum likelihood phylogeny (PhyML) was constructed. A Hierarchical likelihood ratio test (hLRT) with a confidence level of 0.01 suggested that the General Time Reversible (GTR) +G (Rate variation 4 categories) and +T (topology variation) nucleotide substitution model was the most appropriate. 1000 bootstrap replicates were performed with 95% bootstrap branching support cut-off.

## Results

### Presence and assembly of Aedes albopictus densovirus (AalDNV-1) genome in *Ae. aegypti* and *Ae. albopictus* cells

The *Ae. aegypti* cell line Aag2 (Peleg, 1968) is a robust cell line known to be persistently infected with PCLV (Aguiar et al., 2015; Maringer et al., 2017) as well as CFAV (Cammisa-Parks et al., 1992; Scott et al., 2010). In addition to these viruses, a recent study also assembled a contig with the small RNA fraction of Aag2 cells with high pairwise nucleotide similarity to Aedes aegypti densovirus 2 (AeDNV-2) (Aguiar et al., 2015). We conducted total RNA sequencing and qPCR analysis to determine the presence and also the quantity of densovirus genomes in our Aag2 cells.

Total RNA sequencing from Aag2 cells were adapter and quality trimmed resulting in 93,137,517 clean reads for *de novo* assembly. BLASTn analysis of assembled contigs of the non-redundant nucleotide National Center for Biotechnology Information (NCBI) database one 3,937bp contig was identified as 99% identical with highest pairwise nucleotide identity (3680/3721) to Aedes albopictus densovirus (AalDNV-1) (X74945) (Boublik et al., 1994). While this is listed as AalDNV-2 on NCBI, the accepted nomenclature for this densovirus strain is AalDNV-1 (Bergoin and Tijssen, 2010). Assembly was checked by re-mapping to the contig and manually checking for errors. The average coverage of the densovirus contig was 600x (Fig. 1A). Prediction of open reading frames from this contig showed that it is coding complete with no premature stop truncations and represents a *bona fide* DNV genome.

**Figure 1:**
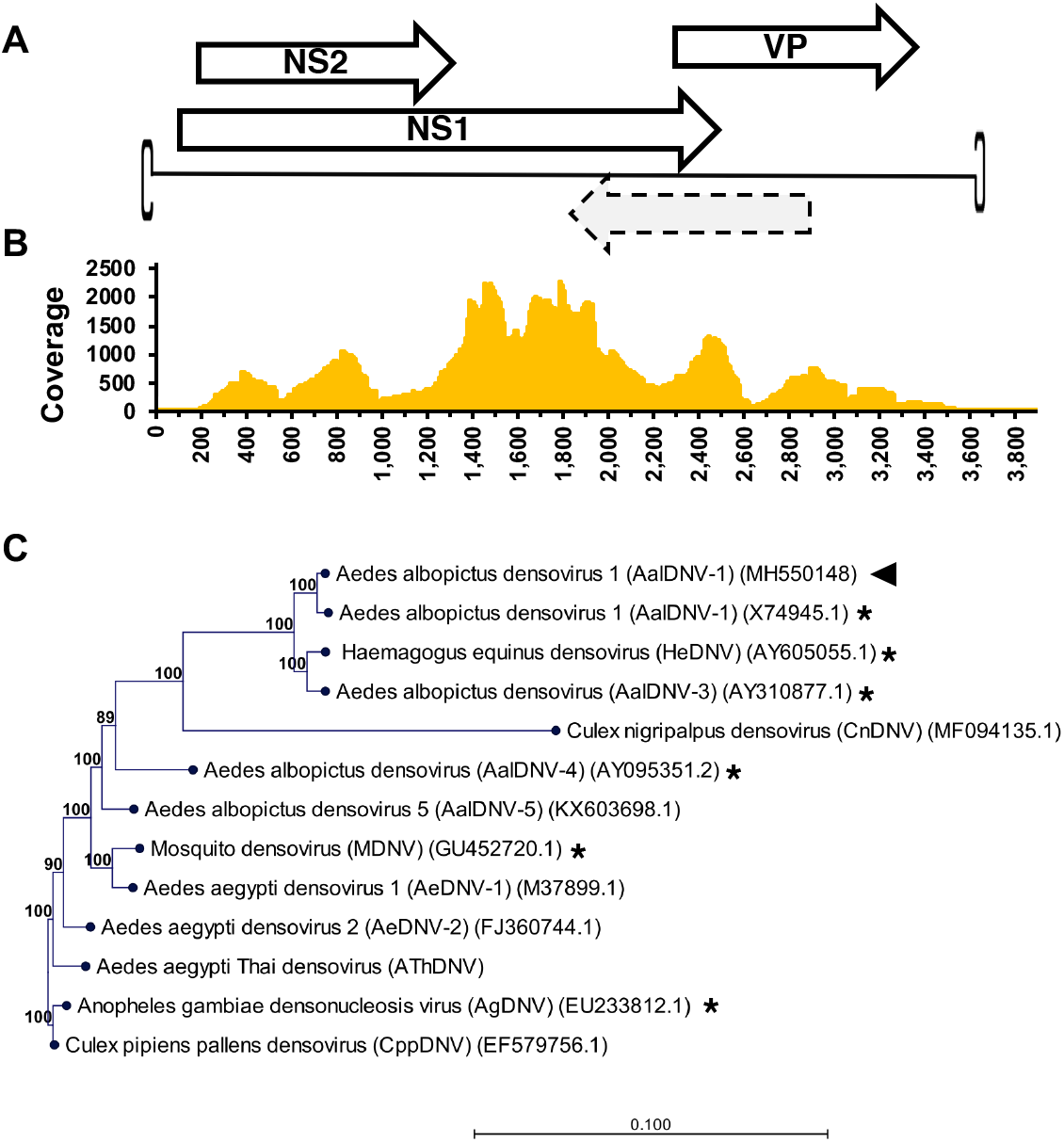
A) The schematic diagram of the genome of Aedes albopictus densovirus (AalDNV-1) assembled from Aag2 cells. B) Mapping coverage of densovirus contig assembled from total RNA. C) Phylogenetic relationship between the AalDNV-1 strain from our Aag2 cells (indicated by an arrowhead) and other densoviruses previously reported in mosquitoes. A star denotes isolation from a mosquito cell line. Maximum likelihood phylogeny (PhyML) between AalDNV-1 strains using a General Time Reversible (GTR) + G +T model with 1000 bootstraps. Branch lengths represent expected numbers of substitutions per nucleotide site.

As we have assembled a related strain to AalDNV-1, and the original designation of this virus, Aedes parvovirus (AaPV) (Boublik et al., 1994), was harvested from *Ae. albopictus* C6/36 cell lines and as these viruses are generally named after host species from which the virus was isolated from, it may not represent the true origin of the virus (Carlson et al., 2006). Phylogenetic analysis of this contig was conducted from published and deposited mosquito DNV strains available in the NCBI nr database: AaeDNV-1 (M37899) (Afanasiev et al., 1991), AaeDNV-2 (AY095351.2) (Chen et al., 2004), AalDNV-1 (X74945) (Boublik et al., 1994), AalDNV-3 (AY310877) AalDNV-5 (KX603698.1), MDNV (BR_07) (Mosimann et al., 2011), Haemagogus equinus densovirus (HeDNV) (AY605055.1) (Paterson et al., 2005), and Aedes Thailand densonucleosis virus (AThDNV) (Kittayapong et al., 1999; Roekring et al., 2006; Zakrzewski et al., 2018). Phylogenetic tree rooted to Anopheles gambiae densovirus (AgDNV) EU233812.1 (Ren et al., 2008) and Culex pipiens pallens densovirus (CpDNV) (EF579756.1) (Zhai et al., 2008) showed that the densovirus persistently infecting Aag2 cells was well supported within a clade from densoviruses previously found in Culicinae cell lines (Fig. 1B). The evolutionary rate between the original Aedes parvovirus genome isolated in 1994 from *Ae. albopictus* C6/36 cells (Boublik et al., 1994) and our contig corresponds to ~5 x 10^-4^ subs/site/year. While this is considered a high substitution rate for a DNA virus, and similar to those of RNA viruses (Jenkins et al., 2002), it has been shown that other mammalian parvoviruses such as canine parvovirus have ∼10^−4^ subs/site/year (Shackelton et al., 2005) and generally ssDNA viruses have rates of evolution generally reported between the range 10^−3^ and 10^−5^ subs/site/year (Jenkins et al., 2002).

### Higher AalDNV-1 infection is associated with *Wolbachia* infection in *Aedes* cells

To test the incidence and abundance of AalDNV-1 in our various *Aedes* cell lines, qPCR was conducted on genomic DNA extracted and purified from *Ae. aegypti* Aag2 cells and different Aag2 derivations: Aag2.*w*MelPop-CLA (hereafter Aag2-Pop) cells, Aag2-Pop cells cleared of *Wolbachia* with tetracycline treatment (hereafter Aag2-PopT). A stably infected Aag2-*w*AlbB line was generated by transinfection of the *Wolbachia w*AlbB strain from Aa23 cells (O’Neill et al., 1997) as well as a tetracycline cured Aag2-*w*AlbBT. Additionally, the *Ae. aegypti* cell line Aa20 was also tested. We also analyzed *Ae. albopictus* cell lines C6/36, Aa23, and its tetracycline cured Aa23-Tet cells. Out of all the cell lines tested, the greatest relative AalDNV-1 copy number relative to host RPS17 was in C6/36 cells (x̄=15,633 n=3) (Fig. 2A). This cell line has been demonstrated to be Dicer-2 deficient and as is used to produce high titres of arboviruses compared with Dicer-2 proficient cell lines (Brackney et al., 2010). Considering the Aag2 cell line and its derivatives, there was a statistically significant increase in AalDNV-1 genome copies between Aag2 (x̄=207.33, n=3) and Aag2-Pop cells (x̄=5,943, n=3) (*p* ≤ 0.01; Student’s t-Test). The tetracycline cleared Aag2-Pop, Aag2-PopT line, had a reduced AalDNV-1 relative genome copy number (x̄=223, n=3) when compared to Aag2-Pop (*p* ≤ 0.05; Student’s t-Test). The *w*AlbB transinfected Aag2 cells Aag2-*w*AlbB had a higher abundance of AalDNV-1 copy number (x̄=877, n=3) compared to its tetracycline cured line Aag2-*w*AlbT (x̄=412, n=3) and also the Aag2 cells (*p* ≤ 0.05; Student’s t-Test). A similar trend was also observed for AalDNV-1 copy number in Aa23 (x̄=0.258, n=3) cells and their tetracycline cured line Aa23-T cells (x̄=0.0684, n=3) (*p* ≤ 0.05; Student’s t-Test).

**Figure 2:**
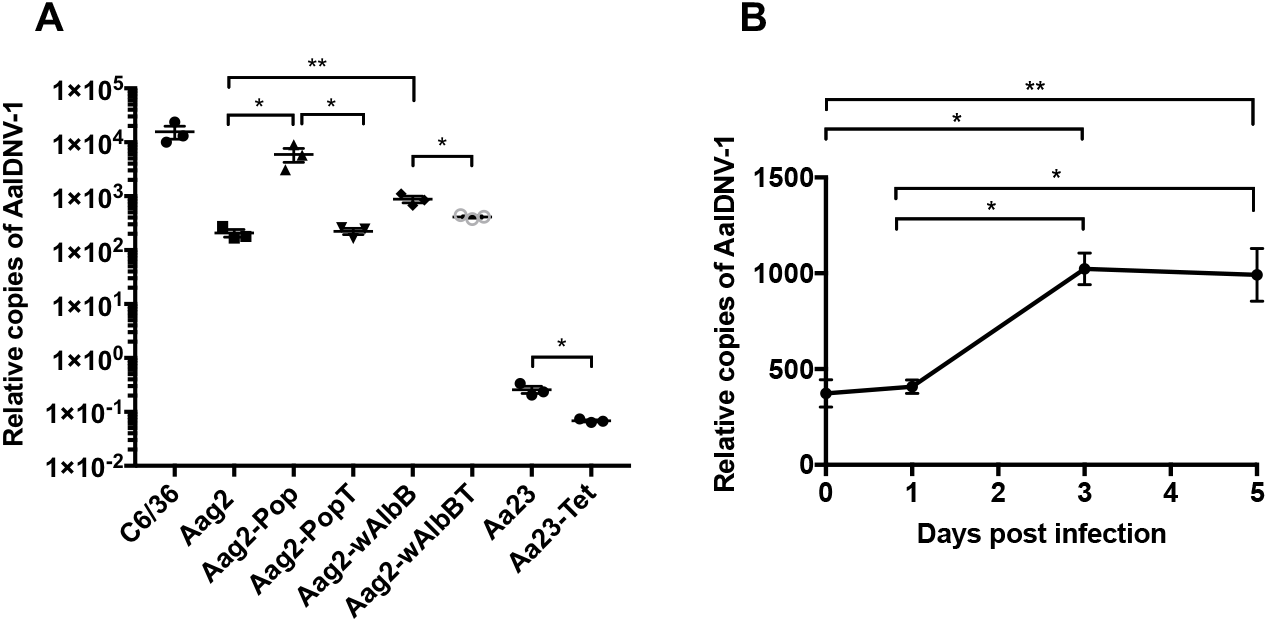
*Ae. aegypti* Aag2 and *Ae. albopictus* Aa23 cells and their derivatives and *Ae. albopictus* C6/36 cells are infected with AalDNV-1 with virus density associated with *Wolbachia* infection. (A) AalDNV-1 levels quantified with qPCR analysis using genomic DNA extracted from Aag2 and Aag2-Pop, cells treated with tetracycline Aag2-PopT, an uninfected control *Ae. aegypti* cell line Aa20 and in *Ae. albopictus* Aa23 cells stably infected with wAlbB. *RPS17* gene was used as a normalizing reference. B) AalDNV-1 particles are infectious. Aa20 cells that lack AalDNV-1 were inoculated with medium from C6/36 cells and collected for DNA extraction at 0, 1, 3 and 5 days post-inoculation. Virus DNA levels, quantified by qPCR, increased over time. * *p* ≤ 0.05, ** *p* ≤ 0.01.

We could not amplify any AalDNV-1 from Aa20 cell line. As such, we used this cell line as a means to find out if the AalDNV-1 particles from both C6/36 and Aag2 cells were infectious. Purified supernatant from both C6/36 (Fig. 2B) and Aag2 medium (data not shown) were shown to increase over a 5-day time course, with Aa20 cells showing higher permissibility to infection and no obvious cytopathic effect.

### AalDNV-1 infection is enhanced by *Wolbachia* in a density dependent manner in *Ae. aegypti* and *Ae. albopictus* cells

To determine the relative density of *Wolbachia* in the aforementioned cells, we conducted qPCR (Fig. 3A) validating the presence of high *Wolbachia* genome copies in Aag2-Pop (x̄=18.9, n=3), Aag2-*w*AlbB (x̄=0.69, n=3) and Aa23 cells (x̄=0.21, n=3) compared with their tetracycline cured lines. To further explore the association between *Wolbachia* presence and higher genome copies of AalDNV-1 in *Aedes* cells, we generated 32 DNA samples from ten successive tetracycline (5*μ*g/mL) treatments of Aag2-Pop and plotted the relationship between density of *Wolbachia* genome copies over genome copies of AalDNV-1. Linear regression analysis showed there was a positive relationship between *Wolbachia* genome copies in these cells (R2= 0.5647; *p* ≤ 0.0001) (Fig. 3B). This trend was also observed when cross checking 28 historical samples from 2017 from nine successive tetracycline (5*μ*g/mL) treatments in the production of the Aag2-Pop-T line (Fig. S1) (R^2^=0.6784; *p* ≤ 0.0001). In addition to tetracycline treatment of Aag2-Pop cells, the stably infected Aag2-*w*AlbB line was monitored every two passages over 30 passages for *Wolbachia* density and AalDNV-1 copy number (n=15). Linear regression analysis showed there was also a positive relationship between *Wolbachia* density (R^2^=0.6779; *p* ≤ 0.001) and AalDNV-1 genome copy number in these cells (Fig. 3C). Taken together, *Wolbachia* appears to enhance AalDNV-1 replication and this enhancement appears to be in a density dependent manner.

**Figure 3:**
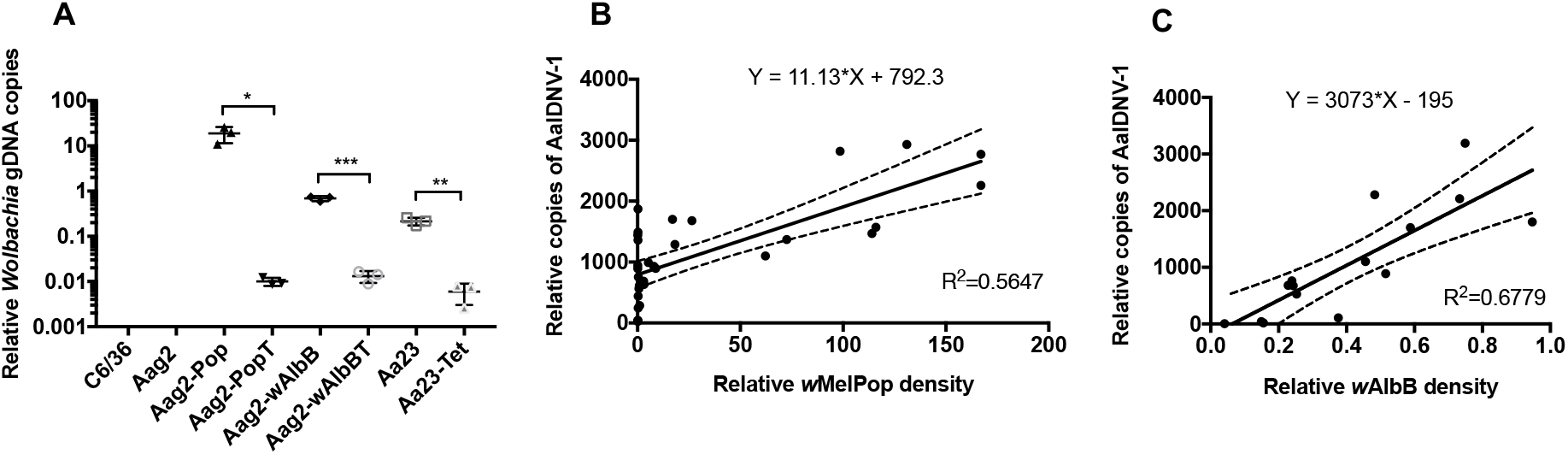
*Wolbachia* enhances AalDNV-1 infection in a density dependent manner. (A) Density of *Wolbachia* in cell lines determined by qPCR using the *wsp* gene relative to the host *RPS17* gene. B) Linear regression analysis of AalDNV-1 genome copies over *Wolbachia* genome copies in Aag2 cells stably infected with *w*MelPop-CLA and sequentially cleared of *Wolbachia* through tetracycline treatment, and C) Aag2 cells transinfected with *w*AlbB over 30 passages. * *p* ≤ 0.05; ** *p* ≤ 0.01; *** *p* ≤ 0.001.

### AalDNV-1 gene transcripts are targeted by the RNAi response

RNAi response is observed in mosquitoes against disparate RNA virus infections. This response includes the microRNA (miRNA) pathway (20-24nt), small interfering RNA (siRNA) pathway (21nt), and the exogenous PIWI-interacting RNAs (piRNA) (25–30 nt) (Blair and Olson, 2015; Hussain et al., 2016; Schnettler et al., 2013). Transcripts of DNA virus genes have been demonstrated to be targeted and modulated by the RNAi pathway in diverse insect DNA virus interactions (Bronkhorst et al., 2012; Jayachandran et al., 2012b; Sabin et al., 2013).

To characterize the RNAi response in Aag2 and Aag2-Pop cells, the small RNA portion of nuclear and cytoplasmic fractions of Aag2 and Aag2-Pop cells previously sequenced was adapter trimmed, pooled and mapped to the assembled AalDNV-1 contig (Mayoral et al., 2014a; Mayoral et al., 2014b). Overall, there was not much difference between the total number of mapped reads between Aag2 (n=28,99; 0.34%) and Aag2-Pop (n=52,693; 0.33%) after normalization of libraries. Additionally, the patterns of regions of viral derived small RNAs mapping to the AalDNV-1 viral genome were the same between both Aag2 (Fig. 4A) and Aag2-Pop cells (Fig. 4B). Total sRNA reads mapped unevenly throughout the AalDNV-1 genome with “hot spot” regions being highly targeted (regions 400-570nt and 2300-2800nt).

**Figure 4:**
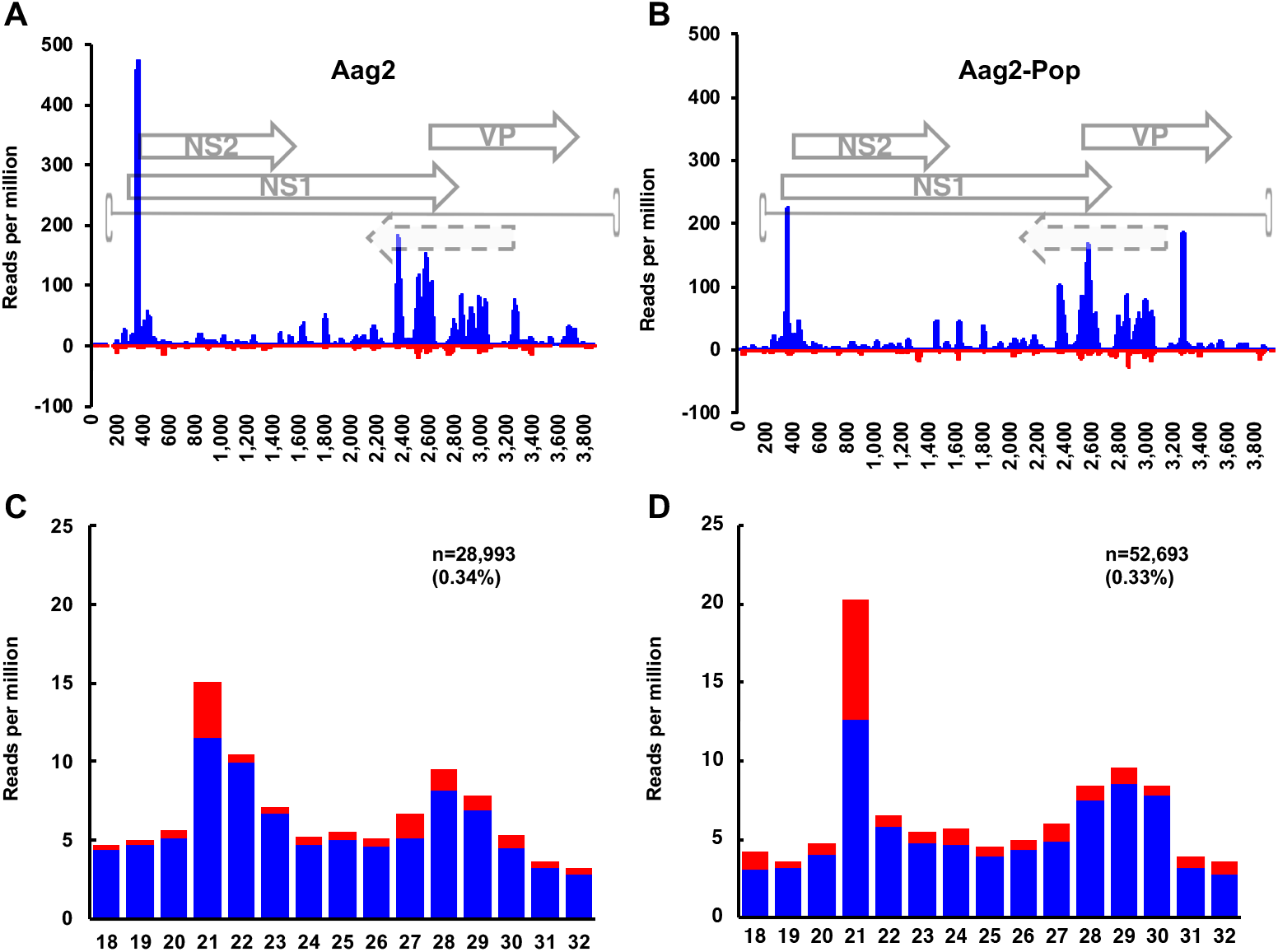
Profile of total normalized 18-32nt reads mapped to the genome strand (blue) and antigenome strand (red) of AalDNV-1 in (A) Aag2 cells (A) and (B) Aag2-Pop cells. Composition and profile of viral derived small RNA reads mapped to AalDNV-1 in (C) Aag2 and (D) Aag2-Pop cells.

In Aag2 cells, the majority of sRNAs mapped to the genome sense (87.45%) orientation than compared with the (12.54%) to the antigenome strand of AalDNV-1. This 7:1 genome:antigenome ratio of mapped reads is similar to previously reported *Culex pipiens molestus* mosquitoes infected with Mosquito densovirus (MDNV) (Ma et al., 2011). In Aag2-Pop, most reads mapped to the genome (80.91%), however interestingly there was a greater number of total reads mapping to the anti-genome (19.08%) 4.24:1 genome:antigenome. Analysis of the composition of sRNA read sizes of AalDNV-1 derived sRNA showed a higher abundance of 21 nt reads (representing viral short interfering RNAs, vsiRNAs) in Aag2-Pop cells (~20% of the total reads) as compared with Aag2 cells (~15%) (Fig. 4C-D). These vsiRNAs are generated by cleavage of viral dsRNA by the RNase III enzyme Dicer-2. This result shows that *Wolbachia* infection of Aag2 cells results in a higher proportion of AalDNV-1 derived vsiRNAs suggesting that either there is greater activity of AalDNV-1 transcription through *Wolbachia* infection or greater genome copies of the virus, or potentially both. Consistently, the qPCR data (Figs. 2 and 3) suggested higher replication of AalDNV-1 in the presence of *Wolbachia*.

While densoviruses do not use dsRNA as a replicative intermediate, it has been previously reported that parvoviruses produce dsRNA (Son et al., 2015; Weber et al., 2006). To further examine the biogenesis of the vsiRNAs in ssDNA virus infection, we trimmed all libraries to 21 nt fractions and re-mapped the reads to the AalDNV-1 genome (Fig. 5A-B). With the exception of high “hot spot” coverage of the genome strand by the sRNAs, we noticed a pattern of “mirroring” of vsiRNA between genome position ~400-3000 nt which corresponds to the *NS1/NS2* transcription and the beginning of the *VP* gene. This pattern of mapped reads has been shown in *Drosophila* cells infected with vaccinia virus (VACV, family *Orthopoxvirus*) and is due to overlapping transcription of genes on both strands of the DNA virus genome (Sabin et al., 2013). This suggests that the biogenesis of these vsiRNAs may be derived through overlapping transcription of genes on the genome and antigenome strands.

**Figure 5:**
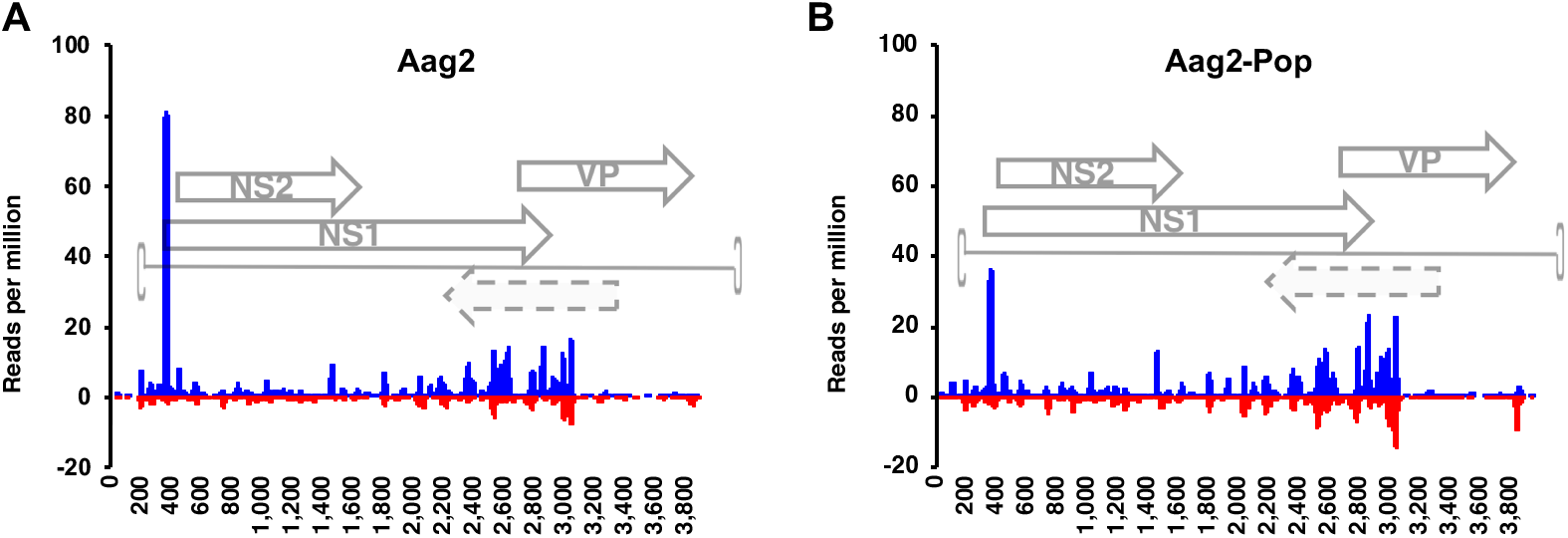
The host RNAi response to AalDNV-1. Mapping profile of the 21nt viral derived siRNAs (vsiRNAs) targeting AalDNV-1 in A) Aag2 and B) Aag2-Pop cells.

The piRNA pathway processes viral RNA into 25-29 nt vpiRNAs. While it has been reported that abundant 25-29 nt sRNA reads were present in *C. pipiens molestus* mosquitoes infected with MDNV (Ma et al., 2011), the vpiRNA signature of these 25-29 nt reads were not examined for the presence of primary and secondary piRNAs. As there is no previous report of the presence of vpiRNAs in the case of DNA viruses of insects, we examined the mapping pattern and also the “ping-pong” signature (U_1_ - A_10_) of AalDNV-1-derived 25-29 nt reads in both Aag2 and Aag2-Pop cells. In comparison to the vsiRNA profile, the vpiRNAs were most abundantly targeted towards the end of the *NS2* and *VP* genes (Fig. 6A-B). Generally, there were also far more vpiRNAs targeting the AalDNV-1 genome in both the pooled datasets (Fig. 4A-B) and individual cytoplasmic and nuclear fractions as compared with the antigenome (Fig. S2).

**Figure 6:**
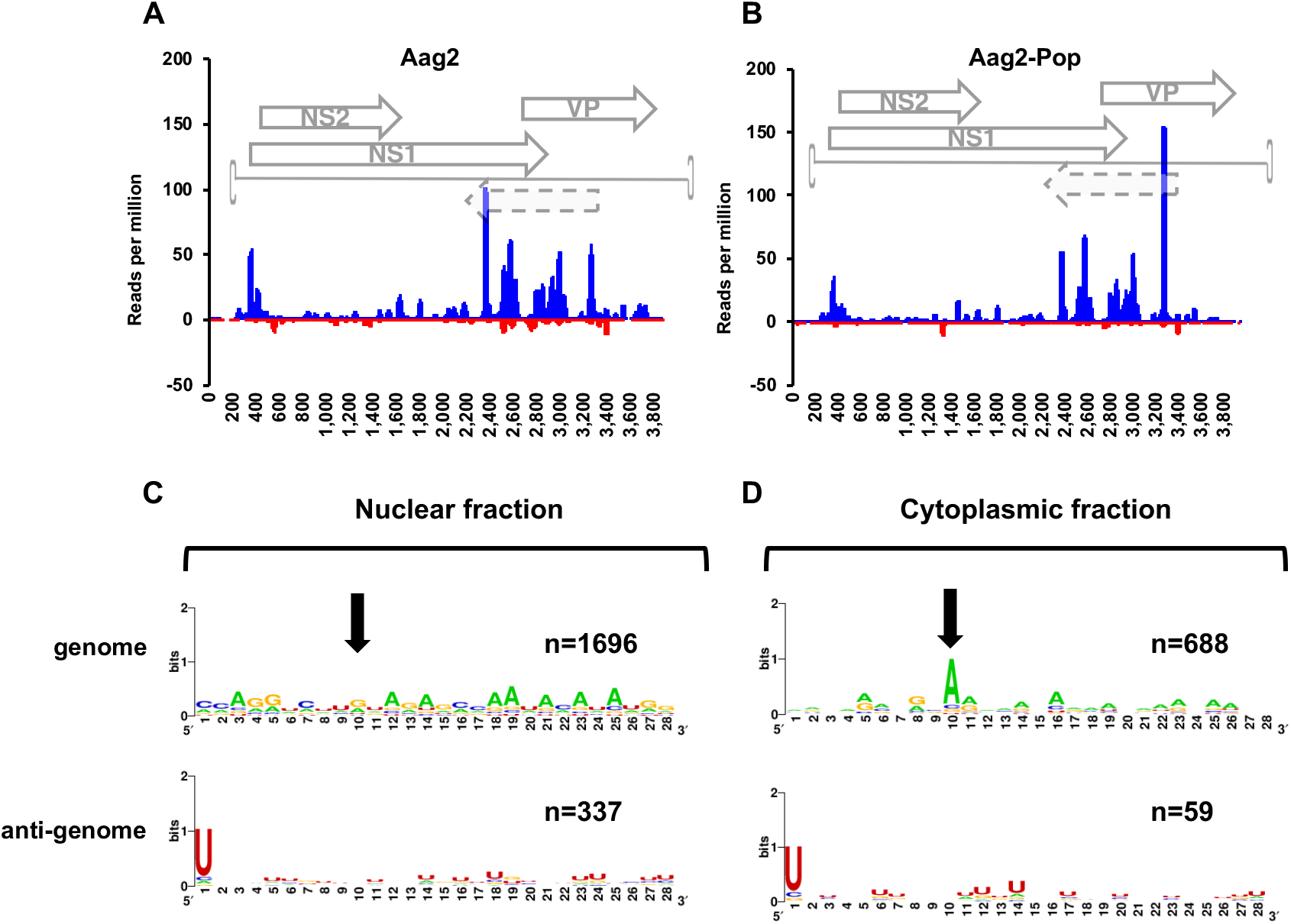
The host piRNA response to AalDNV-1. Mapping profile of 25-29nt viral derived PIWI-interacting piRNAs (vpiRNA) to AalDNV-1 in (A) Aag2 and (B) Aag2-Pop cells. Relative nucleotide frequency and conservation per 28nt vpiRNAs mapped to the AalDNV-1 genome and antigenome in (C) the nuclear and (D) the cytoplasmic fractions of Aag2 cells. Arrow denotes A_10_ position absent in the nuclear fraction. Only Aag2 28nt has been shown as a representative.

In the pooled nuclear and cytoplasmic libraries for both Aag2 and Aag2-Pop cells, we noticed there was a typical “ping-pong” signature (U_1_ - A_10_) of secondary piRNAs in vpiRNAs. However, when we considered just the cytoplasmic and the nuclear fractions of the Aag2 and Aag2-Pop cells individually, we noticed that there was exclusive production of primary piRNAs (U_1_ biased) targeted against the (–) strand and no bias towards an A_10_ signature in the nuclear fraction of both Aag2 and Aag2-Pop cells (Fig. 6C). However, a typical “ping-pong” signature (U_1_ - A_10_) was observed in reads from the cytoplasmic fraction (Fig. 6D).

### A defective AalDNV-1 genome exists in published DNA, RNA and sRNA Aag2 datasets and exclusively produces vsiRNAs

The *Ae. aegypti* Aag2 cell line is a heterogenous embryonic cell line (Peleg, 1968) ubiquitous in arbovirus research laboratories for its robust immune response (Barletta et al., 2012). The Aag2 cell line was recently sequenced through long-read Pac-Bio assembly (Whitfield et al., 2017). To determine if AalDNV-1 was present in the assembly, we scanned the assembled contigs for the presence of AalDNV-1. One 80kb contig (2140) showed a number of hits to our assembled AalDNV-1 genome (Fig. 7A), while it appears that the contig was assembled in the sense and then antisense orientation for ~80kb. Due to the nature of the movie-length PacBIO SMRT Sequencing, if the polymerase runs to the end of the insert, it will continue looping back on the template molecule until the acquisition period ends (Eid et al., 2009). Analysis of the raw data indicated that this was the case as the forward and reverse of the genome were shown immediately in the raw data. Further analysis of the contig showed that it was a partial AalDNV-1 genome assembled by itself, but the *VP* gene of the genome was disrupted whereas the terminal hairpins and *NS1/2* genes remained intact (Fig. 7B).

**Figure 7:**
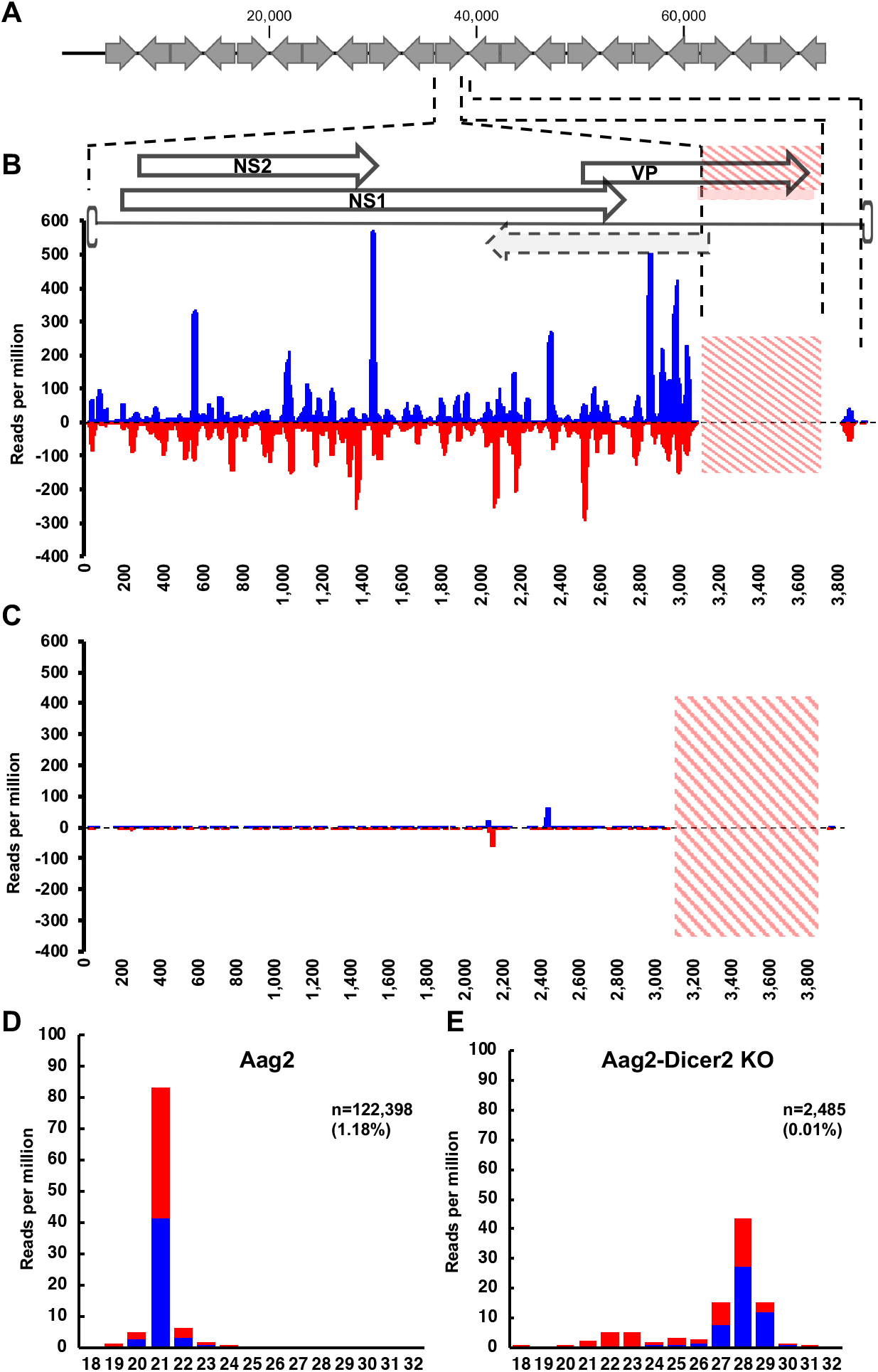
A defective AalDNV-1 particle exists in other Aag2 cells and is targeted almost exclusively by the vsiRNA pathway. A) Contig 2140 from the Aag2 cell line genome project. Arrows denote orientation of the genome. Mapped 18-32nt reads to the AalDNV-1 genome in (B) the parental Aag2 cell line (AF5) and (C) the Dicer-2 knockout Aag2 cells (AF319). Size distribution of mapped sRNA reads from (D) the parental Aag2 cell line (AF5), and E) Dicer-2 knockout (AF319) Aag2 cell line. Red dashed portions in B and C denote the absence of mapped reads to the virus genome.

The Aag2 cell line is RNAi proficient and numerous studies have used the cell line to elucidate components of the siRNA and the piRNA pathways in virus-host interactions. We were interested to re-analyse this collective resource to look at determinants of the RNAi response in Aag2-densovirus interactions. We downloaded small RNA sequencing data from a recent study characterizing the role of the PIWI protein 4 in the antiviral pathway (Varjak et al., 2017). Mapping sRNA reads to our assembled reference contig from the parental Aag2 cell line (AF5) we noticed there was also an absence of mapped reads to the *VP* region of the densovirus genome (Fig. 7B). This was not due to sequencing depth as AalDNV-1 reads comprised as many as 1.08% of the reads (n=122,398). In addition to this absence of mapped reads towards the *VP* gene region, we noticed that vpiRNAs were absent and not produced against this contig (Fig. 7D). Greater than 80% of the mapped reads are 21 nt in size and mapped equally to the genome and antigenome, without the 7:1 genome:antigenome strand bias presented here and previously in mosquitoes infected with DNVs (Ma et al., 2011).

We expanded our search to other Aag2 datasets deposited in the Short Read Archive on NCBI. Subsequent re-mapping of sRNA reads from Aag2 cells indicated that this portion of AalDNV-1 was “silent” or potentially defective in every Aag2 dataset except the total RNA and sRNA data produced from sequencing data from our Aag2 cells. Parvoviruses in mammals have been shown to generate defective particles during high-multiplicity infections (Faust and Ward, 1979). These defective particles vary in size but always retain the terminal palindromes. We suspected that the AalDNV-1 densoviruses common in Aag2 cells may exist as a defective interfering particle. We designed a PCR primer set that encompassed the region that appeared absent in other Aag2 data and were able to amplify an appropriately sized amplicon in our Aag2 cells (data not shown).

In addition to the parental Aag2 (AF5) line, sRNA data from a second Dicer-2 knockout (KO) cell line (AF319) was available for analysis. Reads from the Dicer-2 KO were mapped against our AalDNV-1 reference contig. Once again, there were an absence of reads mapping to the suggested truncated region with only 2,485 reads representing 0.01% of the library (Fig. 7C). In addition, 21 nt vsiRNAs were almost absent against AalDNV-1 in Dicer-2 KO AF319 cells suggesting that vsiRNAs are drastically reduced when Dicer-2 is depleted in the cells (Fig. 7D). Interestingly, the number of vpiRNAs increased in Dicer-2 depleted cells.

As disruption of the *VP* gene results in lack of productive virion production in Aedes densovirus, and that *Ae. aegypti* flanking sequences are absent in the assembled contig from the Aag2 cells used for genome sequencing, the defective AalDNV-1 genome should not be integrated into the host genome and rather extrachromosomally maintained.

## Discussion

### *Wolbachia*-DNA virus interactions

Complex interactions between the endosymbiont *Wolbachia* and its insect host often contribute to modulation of virus titer within the insect. Much of what we currently understand about *Wolbachia*-virus interactions has been characterized using *Drosophila*-*Wolbachia* models and single stranded positive-sense RNA viruses. For example, in *Drosophila simulans* challenged with Flock House virus (FHV, *Nodaviridae*) and Drosophila C virus (DCV, *Dicistroviridae*), it was determined that *Wolbachia* strain was responsible for 93.6% and 70.9% of variance in viral titer, respectively. Further, it was demonstrated that *Wolbachia* density accounted for 86% of *Wolbachia*-mediated protection within the host (Martinez et al., 2017). This density-dependent refractoriness to RNA virus infection conferred by *Wolbachia* has been demonstrated in a number of *in vivo* and *in vitro* studies (Lu et al., 2012; Martinez et al., 2014; Osborne et al., 2012). In *Ae. albopictus* mosquitoes infected with *w*AlbB, clearance of *w*AlbB *in vivo* had no effect on the transmission of DENV or CHIKV but was shown in cells to restrict DENV in a density-dependent manner (Lu et al., 2012). Here, we show that instead of *Wolbachia*-mediated virus refractoriness, in the presence of *Wolbachia* there is a density dependent enhancement of a ssDNA virus. While the exact mechanism of *Wolbachia*-mediated RNA virus restriction has remained elusive, there has been little examination on the mechanism of DNA virus enhancement from *Wolbachia*.

One potential explanation for *Wolbachia*-mediated DNA virus enhancement is in the interplay between reactive oxygen species (ROS) and DNA repair. Increased oxidative stress in various arthropod hosts as a consequence of *Wolbachia* infection is fairly well reported (Brennan et al., 2012; Wong et al., 2015). It is thought that *Wolbachia*-containing vacuoles located in the host cell cytoplasm may act as an extrinsic ROS. Disruption of the redox homeostasis in *Ae. albopictus* Aa23 cells has been shown to increase the incidence of ROS induced DNA damage as demonstrated by increased base 8-oxo-deoxyguanosine in these cells (Brennan et al., 2012). Densoviruses replicate exclusively in the nucleus and require DNA polymerase host machinery for replication. Densoviruses and parvoviruses have also been demonstrated to broadly trigger and require the DNA damage response (DDR) to recruit DNA polymerase co-factors machinery for productive DNA replication (Hristov et al., 2010). It may therefore be possible that the mechanism of *Wolbachia*-mediated DNA virus enhancement lies in the association between ROS induction and the subsequent activation of the DNA damage response (DDR) in *Aedes* cells.

The results reported here expand upon previous findings of *Wolbachia*-DNA viruses interactions in other systems and suggest that both *w*MelPop-CLA, which is a supergroup A *Wolbachia* strain, and *w*AlbB, which is a supergroup B *Wolbachia* strain, can enhance replication of densovirus. As a consequence of this density dependent enhancement of DNA virus replication in insect hosts, it would be advantageous for *Wolbachia* titres to be low enough to have limited effect on the host. *Wolbachia* density has been consistently demonstrated to be significantly lower in the somatic tissues when compared to testes and ovaries of *Drosophila, Ae. albopictus* and transinfected *Ae. aegypti* (Lu et al., 2012; Osborne et al., 2012). Additionally, it has been shown in the transinfected *Ae. aegypti* WB1 strain, *Wolbachia* density is significantly lower in the *Ae. albopictus* HOU strain than in the transinfected *Ae. aegypti* (Lu et al., 2012). We observed a similar trend in our *Ae. aegypti* cells infected with *w*AlbB. The *Ae. aegypti* transinfected cells had a *Wolbachia* density as high as 3-fold more than Aa23 cells.

### Host RNAi response to densovirus

RNAi response, in particular the siRNA pathway, is a powerful anti-viral response against RNA viruses in insects due to the nature of the genome and production of dsRNA intermediates during replication. However, experimental evidence has also shown that the siRNA response is exerted by insects towards dsDNA viruses (Bronkhorst et al., 2012; Jayachandran et al., 2012a; Karamipour et al., 2018; Kemp et al., 2013), most likely against overlapping RNA transcripts produced during transcription of the viral genome. Our results showed for the first time that the host siRNA response could also be triggered by a ssDNA virus. The 21 nt vsiRNAs mostly mapped to ~400-3000 nt which corresponds to the *NS1/NS2* transcription and the beginning of the *VP* gene in AalDNV-1 genome. Interestingly, the amount of 21 nt vsiRNAs mapped to the AalDNV-1 genome increased in *Wolbachia*-infected cells, which correlates with higher replication of the virus in the cells. While it has been previously demonstrated that *Wolbachia w*MelPop-CLA interferes with the localization of Argonaute-1 within *Ae. aegypti* infected cells, there was no evidence for manipulation of the abundance or localization of Argonaute-2 when analysed at the protein level (Hussain et al., 2013b). A recent report showed that in *w*Mel infected Aag2 cells Argonaute-2 transcript levels are significantly upregulated (Terradas et al., 2017) and may contribute in a small way towards *Wolbachia*-mediated blocking of DENV-2 in these cells by enhancing RNAi. However, a more likely reason for the increased vsiRNA production against AalDNV-1 in the presence of *Wolbachia* could be the greater abundance of dsRNAs produced by overlapping transcripts.

As the replication of Aedes albopictus densovirus occurs exclusively in the nucleus, transcripts of genes may be targeted and processed into functional miRNAs from the host Dicer-1 machinery as is reported in other DNA viruses (Hussain and Asgari, 2010; Hussain et al., 2008). As per the previous report of manipulation of the Argonaute-1 protein by *Wolbachia* in *Aedes* cells, it still remains to be elucidated if functional miRNAs are produced by AalDNV-1 and if this is modulated by *Wolbachia* infection.

Production of 25-29 nt vpiRNAs has been shown when insect cells become infected with a number of RNA viruses (Reviewed in (Miesen et al., 2016). We show for the first time that vpiRNAs could also be produced against DNA viruses. A large number of vpiRNAs were found in AalDNV-1 infected Aag2 cells mostly towards the end of the genome. Furthermore, these vpiRNAs demonstrated the characteristic “ping-pong” signature (U_1_ - A_10_), with the A_10_ signature being predominantly found in vpiRNAs in the cytoplasmic fraction. The proteins Piwi5 and Ago3 are the core proteins of the mosquito ping-pong loop and occur almost exclusively in the cytoplasm (Miesen et al., 2015). As the replication of densoviruses are exclusively nuclear, it may be the case that the biogenesis of the primary U_1_ piRNAs in the nucleus may not be from Ago3/Piwi5 cleavage but may in fact be targeted by Zucchini (Zuc).

### Has the AalDNV-1 defective particle cleared AalDNV-1 from most Aag2 cells?

There has been considerable interest in viral derived DNA (vDNA) intermediates produced in mosquitoes in response to RNA virus infection (Goic et al., 2016). Work conducted byPoirier and colleagues showed that defective viral genomes (DVGs) form chimeric forms with host LTR retrotransposases to produce circular vDNA (Poirier et al., 2018). These circular vDNA chimeras were then shown to serve to amplify siRNA-mediated antiviral immunity in insects, as mapping profiles indicated they exclusively produced vsiRNAs (Poirier et al., 2018). Densoviruses are ssDNA viruses until they infect the host. Upon entry into the nucleus, the ssDNA template is “repaired” into a closed circular form. All evidence presented here indicate that the most common and predominant AalDNV-1 in published Aag2 datasets exists as truncated and defective, as it has been demonstrated that plasmids that have disrupted VP genes do replicate and express, but they do not produce virions, and need a “helper” virus with a complete *VP* gene (Afanasiev et al., 1999). The defective particle produces equal genome and antigenome vsiRNAs. In the only other report of RNAi response to DNV infection, there was a 7:1 genome:antigenome bias which are contradictory to other densovirus RNAi reports. The viral profile presented here in other Aag2 cells appears to mirror the exact same profile of the RNAi response that was demonstrated in response to defective vDNA virus genomes as demonstrated previously (Poirier et al., 2018). Also, it seems unlikely that the vDNA was produced from the canonical LTR retrotransposase chimerisation events which has been suggested previously in RNA virus infection (Poirier et al., 2018) and is more likely to be produced through errant DNA genome production.

Interestingly, in Dicer-2 knockout Aag2 cells the 21 nt vsiRNAs basically disappeared but there was an increase in the number of piRNA size sRNAs against the defective genome. Previous research has shown that in the absence of an efficient siRNA response, more piRNA size sRNAs are produced against viruses (Brackney et al., 2010; Scott et al., 2010). However, the number of vpiRNAs produced in Dicer-2 knockout cells is far less than those found in our Aag2 cells. Previous work in *Drosophila* suggest that production of piRNAs against nuclear gene transcripts is controlled by a 100 nt *cis*-regulator elements that triggers piRNA production (Ishizu et al., 2015). The vpiRNAs that are produced in response to AalDNV-1 infection in our non-defective Aag2 cell line are targeted towards the end of the *NS2* and *VP* genes. As the *VP* gene is truncated in the interfering particle, it may be the case that the lack of vpiRNA production is reduced by the absence of a *cis*-regulatory element within the genome.

Amazingly, it appears that this defective interfering particle has managed to predominate in all published Aag2 datasets. The fixation of this defective particle within common Aag2 cells indicates that the vsiRNAs produced from this might have been able to successfully clear the non-defective forms. This suggests that vDNA forms of ssDNA viruses may also work to promote tolerance within mosquitoes to DNA virus infections as is suggested in RNA virus infection.

## Conclusions

Overall, we have shown that in the presence of *Wolbachia*, there is greater levels of AalDNV in *Aedes* cells and this effect is density dependent and could be induced by two different strains of *Wolbachia*. Further, we revealed that both siRNA and piRNA responses are elicited against AalDNV with the corresponding reads increasing in the presence of *Wolbachia*. It has been demonstrated that in *Ae. aegypti* cells infected with mosquito densoviruses different DNV strains demonstrate different pathologies, some inducing cytopathic effect and spontaneous induction of apoptosis (Paterson et al., 2005). As the level of densovirus infection significantly increases in the presence of *Wolbachia,* it may be the case that *Wolbachia*-infected *Aedes* mosquitoes may be more sensitive to DNV pathological effects in the environment. Work presented here also suggests that there is a heterogeneity between laboratory Aag2 cells in regards to the presence of complete or defective AalDNV particles.

## Acknowledgements

The authors would like to acknowledge Dr. Karyn Johnson and Hugo Perdomo Contreras for their valuable insight and discussions. Additionally, we thank Dr. Gareth Price and Dr. Igor Makunin from the University of Queensland for interpretation of the PacBIO sequencing analysis. Funding: This project was funded by the Australian Research Council (grant number DP150101782) to SA and a University of Queensland scholarship to RP.

## Data accessibility Accession numbers

The assembled Aedes albopictus densovirus contig has been deposited in the National Center for Biotechnology Information (NCBI) database under accession number MH550148. Small RNA sequencing data used in this paper is also accessible through GEO series accession number GSE55210.

## Supplementary material

**Figure S1:**
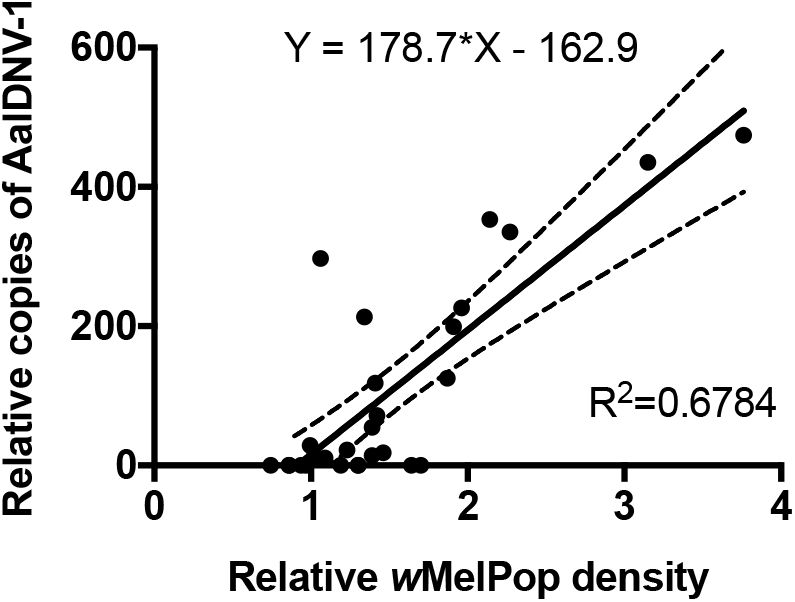
Linear regression analysis of AalDNV-1 genome copies over *Wolbachia* genome copies in Aag2 cells stably infected with *w*MelPop-CLA and sequentially tetracycline cleared using historical 2017 samples.

**Figure S2:**
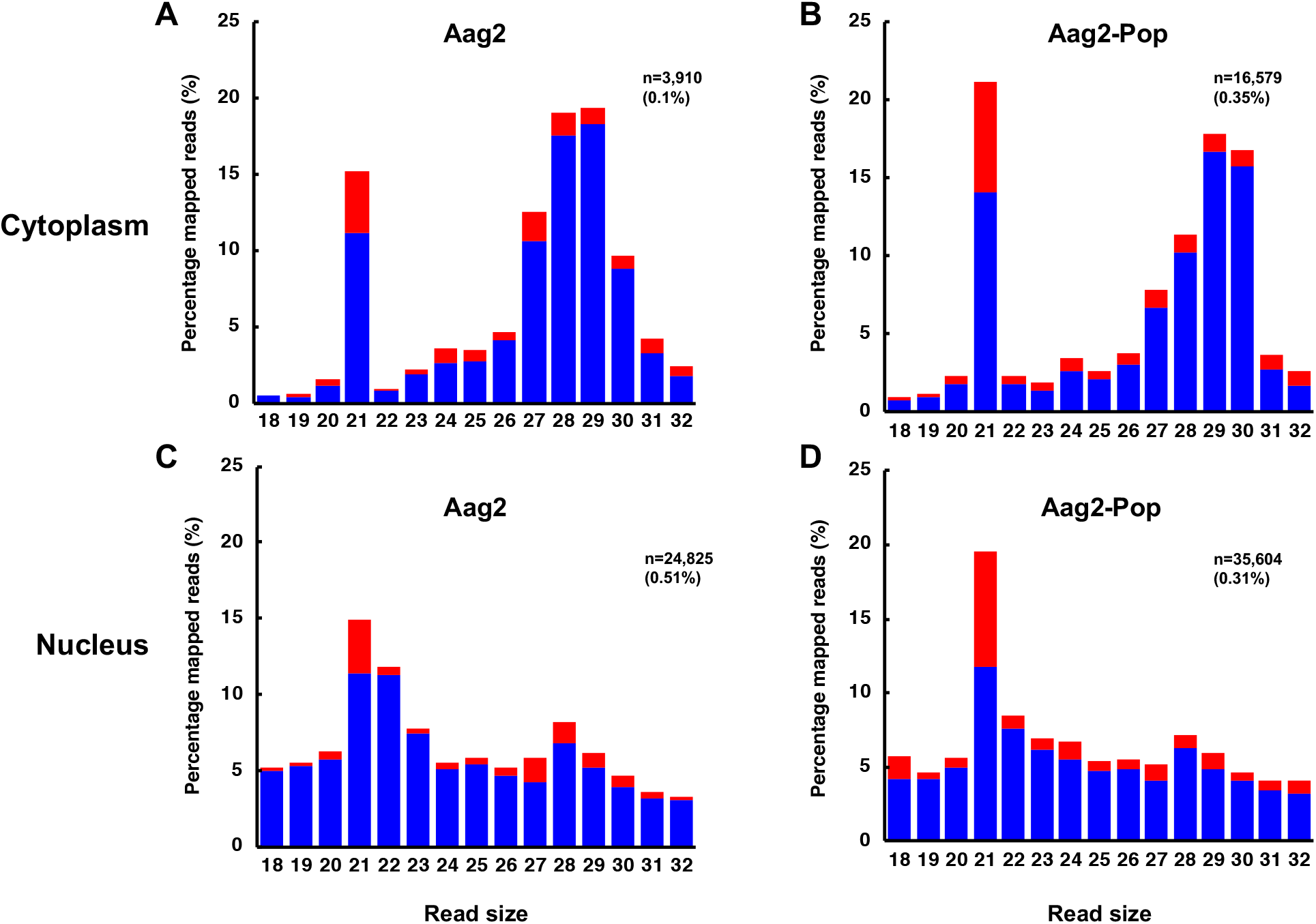
Profile of viral derived small RNA reads mapped to AalDNV-1 in the cytoplasmic fractions of (A) Aag2 and (B) Aag2-Pop cells, and the nuclear fractions of (C) Aag2 and (D) Aag2-Pop cells.

## References

Afanasiev, B.N., Galyov, E.E., Buchatsky, L.P., Kozlov, Y.V., 1991. Nucleotide sequence and genomic organization of Aedes densonucleosis virus. Virology 185, 323–336.

Afanasiev, B.N., Ward, T.W., Beaty, B.J., Carlson, J.O., 1999. Transduction of Aedes aegypti mosquitoes with vectors derived from Aedes densovirus. Virology 257, 62–72.

Aguiar, E.R., Olmo, R.P., Paro, S., Ferreira, F.V., de Faria, I.J., Todjro, Y.M., Lobo, F.P., Kroon, E.G., Meignin, C., Gatherer, D., Imler, J.L., Marques, J.T., 2015. Sequence-independent characterization of viruses based on the pattern of viral small RNAs produced by the host. Nucleic Acids Res 43, 6191–6206.

Aliota, M.T., Peinado, S.A., Velez, I.D., Osorio, J.E., 2016a. The wMel strain of Wolbachia Reduces Transmission of Zika virus by Aedes aegypti. Sci Rep 6, 28792.

Aliota, M.T., Walker, E.C., Uribe Yepes, A., Velez, I.D., Christensen, B.M., Osorio, J.E., 2016b. The wMel Strain of Wolbachia Reduces Transmission of Chikungunya Virus in Aedes aegypti. PLoS Negl Trop Dis 10, e0004677.

Ant, T.H., Herd, C.S., Geoghegan, V., Hoffmann, A.A., Sinkins, S.P., 2018. The Wolbachia strain wAu provides highly efficient virus transmission blocking in Aedes aegypti. PLoS Pathog 14, e1006815.

Asad, S., Parry, R., Asgari, S., 2018. Upregulation of Aedes aegypti Vago1 by Wolbachia and its effect on dengue virus replication. Insect Biochem Mol Biol 92, 45–52.

Barletta, A.B.F., Silva, M.C.L.N., Sorgine, M.H.F., 2012. Validation of Aedes aegypti Aag-2 cells as a model for insect immune studies. Parasites & vectors 5, 148.

Barreau, C., Jousset, F.X., Bergoin, M., 1996. Pathogenicity of the Aedes albopictus parvovirus (AaPV), a denso-like virus, for Aedes aegypti mosquitoes. J Invertebr Pathol 68, 299–309.

Bergoin, M., Tijssen, P., 2010. Densoviruses: a Highly Diverse Group of Arthropod Parvoviruses. Insect Virology, 59–82.

Bhatt, S., Gething, P.W., Brady, O.J., Messina, J.P., Farlow, A.W., Moyes, C.L., Drake, J.M., Brownstein, J.S., Hoen, A.G., Sankoh, O., Myers, M.F., George, D.B., Jaenisch, T., Wint, G.R., Simmons, C.P., Scott, T.W., Farrar, J.J., Hay, S.I., 2013. The global distribution and burden of dengue. Nature 496, 504–507.

Bian, G., Xu, Y., Lu, P., Xie, Y., Xi, Z., 2010. The endosymbiotic bacterium Wolbachia induces resistance to dengue virus in Aedes aegypti. PLoS Pathog 6, e1000833.

Blagrove, M.S., Arias-Goeta, C., Failloux, A.B., Sinkins, S.P., 2012. Wolbachia strain wMel induces cytoplasmic incompatibility and blocks dengue transmission in Aedes albopictus. Proc Natl Acad Sci U S A 109, 255–260.

Blagrove, M.S.C., Arias-Goeta, C., Di Genua, C., Failloux, A.-B., Sinkins, S.P., 2013. A Wolbachia wMel Transinfection in Aedes albopictus Is Not Detrimental to Host Fitness and Inhibits Chikungunya Virus. PLoS Neglected Tropical Diseases 7, e2152.

Blair, C.D., Olson, K.E., 2015. The role of RNA interference (RNAi) in arbovirus-vector interactions. Viruses 7, 820–843.

Boublik, Y., Jousset, F.X., Bergoin, M., 1994. Complete nucleotide sequence and genomic organization of the Aedes albopictus parvovirus (AaPV) pathogenic for Aedes aegypti larvae. Virology 200, 752–763.

Brackney, D.E., Scott, J.C., Sagawa, F., Woodward, J.E., Miller, N.A., Schilkey, F.D., Mudge, J., Wilusz, J., Olson, K.E., Blair, C.D., Ebel, G.D., 2010. C6/36 Aedes albopictus Cells Have a Dysfunctional Antiviral RNA Interference Response. Plos Neglected Tropical Diseases 4.

Brennan, L.J., Haukedal, J.A., Earle, J.C., Keddie, B., Harris, H.L., 2012. Disruption of redox homeostasis leads to oxidative DNA damage in spermatocytes of Wolbachia-infected Drosophila simulans. Insect Mol Biol 21, 510–520.

Bronkhorst, A.W., van Cleef, K.W., Vodovar, N., İnce, İ.A., Blanc, H., Vlak, J.M., Saleh, M.-C., van Rij, R.P., 2012. The DNA virus Invertebrate iridescent virus 6 is a target of the Drosophila RNAi machinery. Proceedings of the National Academy of Sciences 109, E3604–E3613.

Buchatsky, L., Lebedinets, N., Kononko, G., 1997. Densonucleosis of Bloodsucking Mosquitoes: Student Handbook for Biology Majors. Taras Shevchenko’Kiev National University, Kiev.

Cammisa-Parks, H., Cisar, L.A., Kane, A., Stollar, V., 1992. The complete nucleotide sequence of cell fusing agent (CFA): homology between the nonstructural proteins encoded by CFA and the nonstructural proteins encoded by arthropod-borne flaviviruses. Virology 189, 511–524.

Carlson, J., Suchman, E., Buchatsky, L., 2006. Densoviruses for control and genetic manipulation of mosquitoes. Adv Virus Res 68, 361–392.

Chen, S., Cheng, L., Zhang, Q., Lin, W., Lu, X., Brannan, J., Zhou, Z.H., Zhang, J., 2004. Genetic, biochemical, and structural characterization of a new densovirus isolated from a chronically infected Aedes albopictus C6/36 cell line. Virology 318, 123–133.

Coon, K.L., Brown, M.R., Strand, M., 2016. Mosquitoes host communities of bacteria that are essential for development but vary greatly between local habitats. Molecular Ecology 25, 5806–5826.

Crooks, G.E., Hon, G., Chandonia, J.M., Brenner, S.E., 2004. WebLogo: A sequence logo generator. Genome Res 14, 1188–1190.

Dutra, H.L., Rocha, M.N., Dias, F.B., Mansur, S.B., Caragata, E.P., Moreira, L.A., 2016. Wolbachia Blocks Currently Circulating Zika Virus Isolates in Brazilian Aedes aegypti Mosquitoes. Cell Host Microbe 19, 771–774.

Eid, J., Fehr, A., Gray, J., Luong, K., Lyle, J., Otto, G., Peluso, P., Rank, D., Baybayan, P., Bettman, B., Bibillo, A., Bjornson, K., Chaudhuri, B., Christians, F., Cicero, R., Clark, S., Dalal, R., Dewinter, A., Dixon, J., Foquet, M., Gaertner, A., Hardenbol, P., Heiner, C., Hester, K., Holden, D., Kearns, G., Kong, X., Kuse, R., Lacroix, Y., Lin, S., Lundquist, P., Ma, C., Marks, P., Maxham, M., Murphy, D., Park, I., Pham, T., Phillips, M., Roy, J., Sebra, R., Shen, G., Sorenson, J., Tomaney, A., Travers, K., Trulson, M., Vieceli, J., Wegener, J., Wu, D., Yang, A., Zaccarin, D., Zhao, P., Zhong, F., Korlach, J., Turner, S., 2009. Real-time DNA sequencing from single polymerase molecules. Science 323, 133–138.

Faust, E.A., Ward, D.C., 1979. Incomplete genomes of the parvovirus minute virus of mice: selective conservation of genome termini, including the origin for DNA replication. J Virol 32, 276–292.

Frentiu, F.D., Robinson, J., Young, P.R., McGraw, E.A., O’Neill, S.L., 2010. Wolbachia-Mediated Resistance to Dengue Virus Infection and Death at the Cellular Level. Plos One 5.

Gloria-Soria, A., Chiodo, T.G., Powell, J.R., 2018. Lack of Evidence for Natural Wolbachia Infections in Aedes aegypti (Diptera: Culicidae). J Med Entomol.

Goic, B., Stapleford, K.A., Frangeul, L., Doucet, A.J., Gausson, V., Blanc, H., Schemmel-Jofre, N., Cristofari, G., Lambrechts, L., Vignuzzi, M., Saleh, M.C., 2016. Virus-derived DNA drives mosquito vector tolerance to arboviral infection. Nat Commun 7, 12410.

Graham, R.I., Grzywacz, D., Mushobozi, W.L., Wilson, K., 2012. Wolbachia in a major African crop pest increases susceptibility to viral disease rather than protects. Ecol Lett 15, 993–1000.

Hedges, L.M., Brownlie, J.C., O’Neill, S.L., Johnson, K.N., 2008. Wolbachia and virus protection in insects. Science 322, 702.

Hilgenboecker, K., Hammerstein, P., Schlattmann, P., Telschow, A., Werren, J.H., 2008. How many species are infected with Wolbachia? - a statistical analysis of current data. Fems Microbiol Lett 281, 215–220.

Hristov, G., Krämer, M., Li, J., El-Andaloussi, N., Mora, R., Daeffler, L., Zentgraf, H., Rommelaere, J., Marchini, A., 2010. Through Its Nonstructural Protein NS1, Parvovirus H-1 Induces Apoptosis via Accumulation of Reactive Oxygen Species. Journal of Virology 84, 5909–5922.

Hussain, M., Asgari, S., 2010. Functional analysis of a cellular microRNA in insect host-ascovirus interaction. J Virol 84, 612–620.

Hussain, M., Etebari, K., Asgari, S., 2016. Chapter Seven - Functions of Small RNAs in Mosquitoes, in: Raikhel, A.S. (Ed.), Advances in Insect Physiology. Academic Press, pp. 189–222.

Hussain, M., Lu, G., Torres, S., Edmonds, J.H., Kay, B.H., Khromykh, A.A., Asgari, S., 2013a. Effect of Wolbachia on replication of West Nile virus in a mosquito cell line and adult mosquitoes. J Virol 87, 851–858.

Hussain, M., O’Neill, S.L., Asgari, S., 2013b. Wolbachia interferes with the intracellular distribution of Argonaute 1 in the dengue vector Aedes aegypti by manipulating the host microRNAs. RNA Biol 10, 1868–1875.

Hussain, M., Taft, R.J., Asgari, S., 2008. An insect virus-encoded microRNA regulates viral replication. J Virol 82, 9164–9170.

Ishizu, H., Iwasaki, Y.W., Hirakata, S., Ozaki, H., Iwasaki, W., Siomi, H., Siomi, M.C., 2015. Somatic Primary piRNA Biogenesis Driven by cis-Acting RNA Elements and trans-Acting Yb. Cell reports 12, 429–440.

Iturbe-Ormaetxe, I., Woolfit, M., Rances, E., Duplouy, A., O’Neill, S.L., 2011. A simple protocol to obtain highly pure Wolbachia endosymbiont DNA for genome sequencing. J Microbiol Methods 84, 134–136.

Jayachandran, B., Hussain, M., Asgari, S., 2012a. RNA interference as a cellular defense mechanism against the DNA virus baculovirus. J Virol 86, 13729–13734.

Jayachandran, B., Hussain, M., Asgari, S., 2012b. RNA interference as a cellular defense mechanism against the DNA virus baculovirus. Journal of virology 86, 13729–13734.

Jenkins, G.M., Rambaut, A., Pybus, O.G., Holmes, E.C., 2002. Rates of molecular evolution in RNA viruses: a quantitative phylogenetic analysis. J Mol Evol 54, 156–165.

Jeyaprakash, A., Hoy, M.A., 2000. Long PCR improves Wolbachia DNA amplification: wsp sequences found in 76% of sixty-three arthropod species. Insect Mol Biol 9, 393–405.

Johnson, R.M., Rasgon, J.L., 2018. Densonucleosis viruses (‘densoviruses’) for mosquito and pathogen control. Current Opinion in Insect Science 28, 90–97.

Joubert, D.A., O’Neill, S.L., 2017. Comparison of Stable and Transient Wolbachia Infection Models in Aedes aegypti to Block Dengue and West Nile Viruses. PLoS Negl Trop Dis 11, e0005275.

Joubert, D.A., Walker, T., Carrington, L.B., De Bruyne, J.T., Kien, D.H., Hoang Nle, T., Chau, N.V., Iturbe-Ormaetxe, I., Simmons, C.P., O’Neill, S.L., 2016. Establishment of a Wolbachia Superinfection in Aedes aegypti Mosquitoes as a Potential Approach for Future Resistance Management. PLoS Pathog 12, e1005434.

Jousset, F.X., Barreau, C., Boublik, Y., Cornet, M., 1993. A parvo-like virus persistently infecting a C6/36 clone of Aedes albopictus mosquito cell line and pathogenic for Aedes aegypti larvae. Virus Res 29, 99–114.

Juliano, S.A., Lounibos, L.P., 2005. Ecology of invasive mosquitoes: effects on resident species and on human health. Ecol Lett 8, 558–574.

Karamipour, N., Fathipour, Y., Talebi, A.A., Asgari, S., Mehrabadi, M., 2018. Small interfering RNA pathway contributes to antiviral immunity in Spodoptera frugiperda (Sf9) cells following Autographa californica multiple nucleopolyhedrovirus infection. Insect Biochem Mol Biol 101, 24–31.

Kemp, C., Mueller, S., Goto, A., Barbier, V., Paro, S., Bonnay, F., Dostert, C., Troxler, L., Hetru, C., Meignin, C., Pfeffer, S., Hoffmann, J.A., Imler, J.L., 2013. Broad RNA interference-mediated antiviral immunity and virus-specific inducible responses in Drosophila. J Immunol 190, 650–658.

Kittayapong, P., Baisley, K.J., O’Neill, S.L., 1999. A mosquito densovirus infecting Aedes aegypti and Aedes albopictus from Thailand. Am J Trop Med Hyg 61, 612–617.

Ledermann, J.P., Suchman, E.L., Black, W.C.t., Carlson, J.O., 2004. Infection and pathogenicity of the mosquito densoviruses AeDNV, HeDNV, and APeDNV in Aedes aegypti mosquitoes (Diptera: Culicidae). J Econ Entomol 97, 1828–1835.

Lu, P., Bian, G., Pan, X., Xi, Z., 2012. Wolbachia induces density-dependent inhibition to dengue virus in mosquito cells. PLoS neglected tropical diseases 6, e1754.

Ma, M., Huang, Y., Gong, Z., Zhuang, L., Li, C., Yang, H., Tong, Y., Liu, W., Cao, W., 2011. Discovery of DNA viruses in wild-caught mosquitoes using small RNA high throughput sequencing. PLoS One 6, e24758.

Maringer, K., Yousuf, A., Heesom, K.J., Fan, J., Lee, D., Fernandez-Sesma, A., Bessant, C., Matthews, D.A., Davidson, A.D., 2017. Proteomics informed by transcriptomics for characterising active transposable elements and genome annotation in Aedes aegypti. BMC Genomics 18, 101.

Martinez, J., Longdon, B., Bauer, S., Chan, Y.S., Miller, W.J., Bourtzis, K., Teixeira, L., Jiggins, F.M., 2014. Symbionts commonly provide broad spectrum resistance to viruses in insects: a comparative analysis of Wolbachia strains. PLoS Pathog 10, e1004369.

Martinez, J., Tolosana, I., Ok, S., Smith, S., Snoeck, K., Day, J.P., Jiggins, F.M., 2017. Symbiont strain is the main determinant of variation in Wolbachia-mediated protection against viruses across Drosophila species. Mol Ecol 26, 4072–4084.

Mayoral, J.G., Etebari, K., Hussain, M., Khromykh, A.A., Asgari, S., 2014a. Wolbachia infection modifies the profile, shuttling and structure of microRNAs in a mosquito cell line. PLoS One 9, e96107.

Mayoral, J.G., Hussain, M., Joubert, D.A., Iturbe-Ormaetxe, I., O’Neill, S.L., Asgari, S., 2014b. Wolbachia small noncoding RNAs and their role in cross-kingdom communications. Proc Natl Acad Sci U S A 111, 18721–18726.

Miesen, P., Girardi, E., van Rij, R.P., 2015. Distinct sets of PIWI proteins produce arbovirus and transposon-derived piRNAs in Aedes aegypti mosquito cells. Nucleic Acids Res 43, 6545–6556.

Miesen, P., Joosten, J., van Rij, R.P., 2016. PIWIs Go Viral: Arbovirus-Derived piRNAs in Vector Mosquitoes. PLoS Pathog 12, e1006017.

Moreira, L.A., Iturbe-Ormaetxe, I., Jeffery, J.A., Lu, G., Pyke, A.T., Hedges, L.M., Rocha, B.C., Hall-Mendelin, S., Day, A., Riegler, M., Hugo, L.E., Johnson, K.N., Kay, B.H., McGraw, E.A., van den Hurk, A.F., Ryan, P.A., O’Neill, S.L., 2009. A Wolbachia symbiont in Aedes aegypti limits infection with dengue, Chikungunya, and Plasmodium. Cell 139, 1268–1278.

Mosimann, A.L.P., Bordignon, J., Mazzarotto, G.C.A., Motta, M.C.M., Hoffmann, F., Santos, C.N.D.d., 2011. Genetic and biological characterization of a densovirus isolate that affects dengue virus infection. Memórias do Instituto Oswaldo Cruz 106, 285–292.

O’Neill, S.L., Pettigrew, M.M., Sinkins, S.P., Braig, H.R., Andreadis, T.G., Tesh, R.B., 1997. In vitro cultivation of Wolbachia pipientis in an Aedes albopictus cell line. Insect Mol Biol 6, 33–39.

O’Neill, S.L., Kittayapong, P., Braig, H.R., Andreadis, T.G., Gonzalez, J.P., Tesh, R.B., 1995. Insect Densoviruses May Be Widespread in Mosquito Cell-Lines. Journal of General Virology 76, 2067–2074.

Osborne, S.E., Iturbe-Ormaetxe, I., Brownlie, J.C., O’Neill, S.L., Johnson, K.N., 2012. Antiviral protection and the importance of Wolbachia density and tissue tropism in Drosophila simulans. Appl Environ Microbiol 78, 6922–6929.

Parry, R., Asgari, S., 2018. Aedes anphevirus (AeAV): an insect-specific virus distributed worldwide in Aedes aegypti mosquitoes that has complex interplays with Wolbachia and dengue virus infection in cells. J Virol.

Paterson, A., Robinson, E., Suchman, E., Afanasiev, B., Carlson, J., 2005. Mosquito densonucleosis viruses cause dramatically different infection phenotypes in the C6/36 Aedes albopictus cell line. Virology 337, 253–261.

Peleg, J., 1968. Growth of arboviruses in monolayers from subcultured mosquito embryo cells. Virology 35, 617–619.

Poirier, E.Z., Goic, B., Tome-Poderti, L., Frangeul, L., Boussier, J., Gausson, V., Blanc, H., Vallet, T., Loyd, H., Levi, L.I., Lanciano, S., Baron, C., Merkling, S.H., Lambrechts, L., Mirouze, M., Carpenter, S., Vignuzzi, M., Saleh, M.C., 2018. Dicer-2-Dependent Generation of Viral DNA from Defective Genomes of RNA Viruses Modulates Antiviral Immunity in Insects. Cell Host Microbe 23, 353–365 e358.

Pudney, M., Varma, M., Leake, C., 1979. Establishment of cell lines from larvae of culicine (Aedes species) and anopheline mosquitoes. TCA manual/Tissue Culture Association 5, 997–1002.

Ren, X., Hoiczyk, E., Rasgon, J.L., 2008. Viral paratransgenesis in the malaria vector Anopheles gambiae. PLoS pathogens 4, e1000135.

Ridley, A.W., Hereward, J.P., Daglish, G.J., Raghu, S., McCulloch, G.A., Walter, G.H., 2016. Flight of Rhyzopertha dominica (Coleoptera: Bostrichidae)-a Spatio-Temporal Analysis With Pheromone Trapping and Population Genetics. J Econ Entomol 109, 2561–2571.

Roekring, S., Flegel, T.W., Malasit, P., Kittayapong, P., 2006. Challenging successive mosquito generations with a densonucleosis virus yields progressive survival improvement but persistent, innocuous infections. Developmental & Comparative Immunology 30, 878–892.

Sabin, L.R., Zheng, Q., Thekkat, P., Yang, J., Hannon, G.J., Gregory, B.D., Tudor, M., Cherry, S., 2013. Dicer-2 processes diverse viral RNA species. PloS one 8, e55458.

Schnettler, E., Donald, C.L., Human, S., Watson, M., Siu, R.W., McFarlane, M., Fazakerley, J.K., Kohl, A., Fragkoudis, R., 2013. Knockdown of piRNA pathway proteins results in enhanced Semliki Forest virus production in mosquito cells. J Gen Virol 94, 1680–1689.

Schnettler, E., Sreenu, V.B., Mottram, T., McFarlane, M., 2016. Wolbachia restricts insect-specific flavivirus infection in Aedes aegypti cells. J Gen Virol 97, 3024–3029.

Scott, J.C., Brackney, D.E., Campbell, C.L., Bondu-Hawkins, V., Hjelle, B., Ebel, G.D., Olson, K.E., Blair, C.D., 2010. Comparison of dengue virus type 2-specific small RNAs from RNA interference-competent and -incompetent mosquito cells. PLoS Negl Trop Dis 4, e848.

Shackelton, L.A., Parrish, C.R., Truyen, U., Holmes, E.C., 2005. High rate of viral evolution associated with the emergence of carnivore parvovirus. Proc Natl Acad Sci U S A 102, 379–384.

Singh, K.R.P., 1967. Cell cultures derived from larvae of Aedes albopictus (Skuse) AND Aedes aegypti (L.). Current Science 36, 506–508.

Sinkins, S.P., Braig, H.R., O’Neill, S.L., 1995. Wolbachia superinfections and the expression of cytoplasmic incompatibility. Proc Biol Sci 261, 325–330.

Son, K.N., Liang, Z., Lipton, H.L., 2015. Double-Stranded RNA Is Detected by Immunofluorescence Analysis in RNA and DNA Virus Infections, Including Those by Negative-Stranded RNA Viruses. J Virol 89, 9383–9392.

Teixeira, L., Ferreira, A., Ashburner, M., 2008. The bacterial symbiont Wolbachia induces resistance to RNA viral infections in Drosophila melanogaster. PLoS Biol 6, e2.

Terradas, G., Joubert, D.A., McGraw, E.A., 2017. The RNAi pathway plays a small part in Wolbachia-mediated blocking of dengue virus in mosquito cells. Scientific Reports 7.

van den Hurk, A.F., Hall-Mendelin, S., Pyke, A.T., Frentiu, F.D., McElroy, K., Day, A., Higgs, S., O’Neill, S.L., 2012. Impact of Wolbachia on infection with chikungunya and yellow fever viruses in the mosquito vector Aedes aegypti. PLoS Negl Trop Dis 6, e1892.

Varjak, M., Maringer, K., Watson, M., Sreenu, V.B., Fredericks, A.C., Pondeville, E., Donald, C.L., Sterk, J., Kean, J., Vazeille, M., Failloux, A.B., Kohl, A., Schnettler, E., 2017. Aedes aegypti Piwi4 Is a Noncanonical PIWI Protein Involved in Antiviral Responses. mSphere 2.

Walker, T., Johnson, P.H., Moreira, L.A., Iturbe-Ormaetxe, I., Frentiu, F.D., McMeniman, C.J., Leong, Y.S., Dong, Y., Axford, J., Kriesner, P., Lloyd, A.L., Ritchie, S.A., O’Neill, S.L., Hoffmann, A.A., 2011. The wMel Wolbachia strain blocks dengue and invades caged Aedes aegypti populations. Nature 476, 450–453.

Weber, F., Wagner, V., Rasmussen, S.B., Hartmann, R., Paludan, S.R., 2006. Double-stranded RNA is produced by positive-strand RNA viruses and DNA viruses but not in detectable amounts by negative-strand RNA viruses. J Virol 80, 5059–5064.

Whitfield, Z.J., Dolan, P.T., Kunitomi, M., Tassetto, M., Seetin, M.G., Oh, S., Heiner, C., Paxinos, E., Andino, R., 2017. The Diversity, Structure, and Function of Heritable Adaptive Immunity Sequences in the Aedes aegypti Genome. Current Biology 27, 3511–3519. e3517.

Wong, Z.S., Brownlie, J.C., Johnson, K.N., 2015. Oxidative stress correlates with Wolbachia-mediated antiviral protection in Wolbachia-Drosophila associations. Appl Environ Microbiol 81, 3001–3005.

Ye, Y.H., Carrasco, A.M., Frentiu, F.D., Chenoweth, S.F., Beebe, N.W., van den Hurk, A.F., Simmons, C.P., O’Neill, S.L., McGraw, E.A., 2015. Wolbachia Reduces the Transmission Potential of Dengue-Infected Aedes aegypti. PLoS Negl Trop Dis 9, e0003894.

Zakrzewski, M., Rašić, G., Darbro, J., Krause, L., Poo, Y.S., Filipović, I., Parry, R., Asgari, S., Devine, G., Suhrbier, A., 2018. Mapping the virome in wild-caught Aedes aegypti from Cairns and Bangkok. Scientific reports 8, 4690.

Zhai, Y.-g., Lv, X.-j., Sun, X.-h., Fu, S.-h., Fen, Y., Tong, S.-x., Wang, Z.-x., Tang, Q., Attoui, H., Liang, G.-d., 2008. Isolation and characterization of the full coding sequence of a novel densovirus from the mosquito Culex pipiens pallens. Journal of General Virology 89, 195–199.

Zhang, G., Etebari, K., Asgari, S., 2016. Wolbachia suppresses cell fusing agent virus in mosquito cells. J Gen Virol 97, 3427–3432.

Zhou, W., Rousset, F., O’Neil, S., 1998. Phylogeny and PCR-based classification of Wolbachia strains using wsp gene sequences. Proc Biol Sci 265, 509–515.

